# Quorum sensing modulatory effect of sound stimulation on *Serratia marcescens* and *Pseudomonas aeruginosa*

**DOI:** 10.1101/072850

**Authors:** Vijay Kothari, Pooja Patel, Chinmayi Joshi, Brijesh Mishra, Shashikant Dubey, Milan Mehta

## Abstract

Effect of nine different mono-frequency sound stimuli on two gram-negative bacteria (*Pseudomonas aeruginosa* and *Serratia marcescens*) was investigated. Frequency of the test sound ranged from 100 Hz to 2,000 Hz. Both the test bacteria responded differently to sonic stimulation. Sound corresponding to 600 HZ caused a notable reduction in quorum sensing (QS) regulated production of the pigment pyoverdine by *P. aeruginosa*. 400 Hz sound affected prodigiosin production by *S. marcescens* the most. 500 Hz sound could enhance prodigiosin production without affecting growth of the producing bacterium, suggesting the effect purely to be QS modulatory. This study has demonstrated the capacity of the sound waves of affecting bacterial growth and quorum sensing regulated metabolite production.

## Introduction

Sound is one of the most widely distributed environmental factors. One or another kind of sound can be found in almost all corners of the natural world. Effect of a variety of environmental factors such as temperature, light, pH, oxygen concentration, etc. on microorganisms has been studied well. However despite sound being a universally present factor, its effect on the microbial life forms has not received enough attention from the community of microbiologists. Though microorganisms do not possess any auditory cell component, there are reports (Ying *et al*., 2009; Sarvaiya and Kothari, 2015b; Kim, 2016) indicating them to be affected notably by sonic stimulation. The precise mechanisms explaining how microbes sense and respond to external sound stimuli are not yet understood. It is required to conduct the investigations regarding influence of sound on many different bacteria, algae, protozoa, and fungi, to find out whether there is any common pattern underlying microbial response to sound. Ultrasound in general is known to be deleterious to microbial cells (Ying *et al*., 2009), and is widely used in cell lysis protocols. However there is no large body of literature to throw light on interaction of sonic range (20 Hz - 20 KHz) of sound with microbes.

Our previous studies (Kothari and Sarvaiya, 2015b; Shah *et al*., 2016) indicated *Serratia marcescens* responding differently to sound stimulation as compared to other test organisms. In the present study, we subjected *S. marcescens* and *Pseudomonas aeruginosa* to sonic stimulation at different sound frequencies, and observed the effect on their growth and quorum sensing (QS) regulated pigment production. In our previously published studies, we used multi-frequency sound in form of music, whereas in the present study mono-frequency sound was employed to investigate whether these two bacteria respond to sonic stimulation in a frequency dependent fashion.

## Materials and Methods

### Bacterial culture

*Serratia marcescens* (MTCC 97) was procured from the Microbial Type Culture Collection (MTCC), Chandigarh. This organism was grown in nutrient broth (HiMedia, Mumbai) supplemented with 1% v/v glycerol (HiMedia,). Incubation was made at 28° C for 48 h. *Pseudomonas aeruginosa* was procured from Microbiology Department of M.G. Science Institute, Ahmedabad. The organism was grown in *Pseudomonas* broth (2% peptone; 1% potassium sulphate; 0.14% MgCl_2_; pH 7±0.2). Incubation was made at 35°C for 24 h.

### Sound generation

Sound beep (s) of required frequency was generated using NCH^®^ tone generator. The sound file played during the experiment was prepared using WavePad Sound Editor Masters Edition v.5.5 in such a way that there is a time gap of one second between two consecutive beep sounds.

### Sound stimulation

Inoculum of the test bacterium was prepared from its activated culture, in sterile normal saline. Optical density of this inoculum was adjusted to 0.08-0.10 at 625 nm (Agilent Technologies Cary 60 UV-vis spectrophotometer). The test tubes (Borosil, 25x100 mm; 38 mL) containing inoculated growth medium (6 mL including 5%v/v inoculum) were put into a glass chamber (Actira, L: 250 x W: 250 x H: 150 mm). A speaker (Minix sound bar, Maxtone Electronics Pvt. Ltd., Thane) was put in this glass chamber at the distance of 15 cm from the inoculated test tubes. Sound delivery from the speaker was provided throughout the period of incubation. This glass chamber was covered with a glass lid, and one layer of loose-fill shock absorber polystyrene, in such a way that the polystyrene layer gets placed below the glass lid. Silicone grease was applied on the periphery of the glass chamber coming in contact with the polystyrene material. This type of packaging was done to minimize any possible leakage of sound from inside of the chamber, and also to avoid any interference from external sound. Similar chamber was used to house the ‘control’ (i.e. not subjected to sound stimulation) group test tubes. One non-playing speaker was also placed in the glass chamber used for the control tubes at a distance of 15 cm from tubes, where no electricity was supplied and no sound was generated (Kothari et al., 2016). Intensity of sound, measured with a sound level meter (acd machine control Ltd.) at a distance of 15 cm from the speaker, varied with the frequency, in the range of 75-99 dB (Table 1). Sound level in control chamber was found to be below the detection level of the sound level meter. Schematic of the whole experimental set-up is shown in Figure 1. Intermittent mixing of the contents of the test tubes to minimize heterogeneity was achieved by vortexing the tubes at an interval of every 3 h using a vortex mixer. Whenever the tubes were taken out for vortexing, each time positions of tubes of a single chamber were inter-changed, and their direction with respect to speaker was changed by rotating them 180°. This was done to achieve a high probability of almost equal sound exposure to all the tubes, and their content.

**Figure 1.**
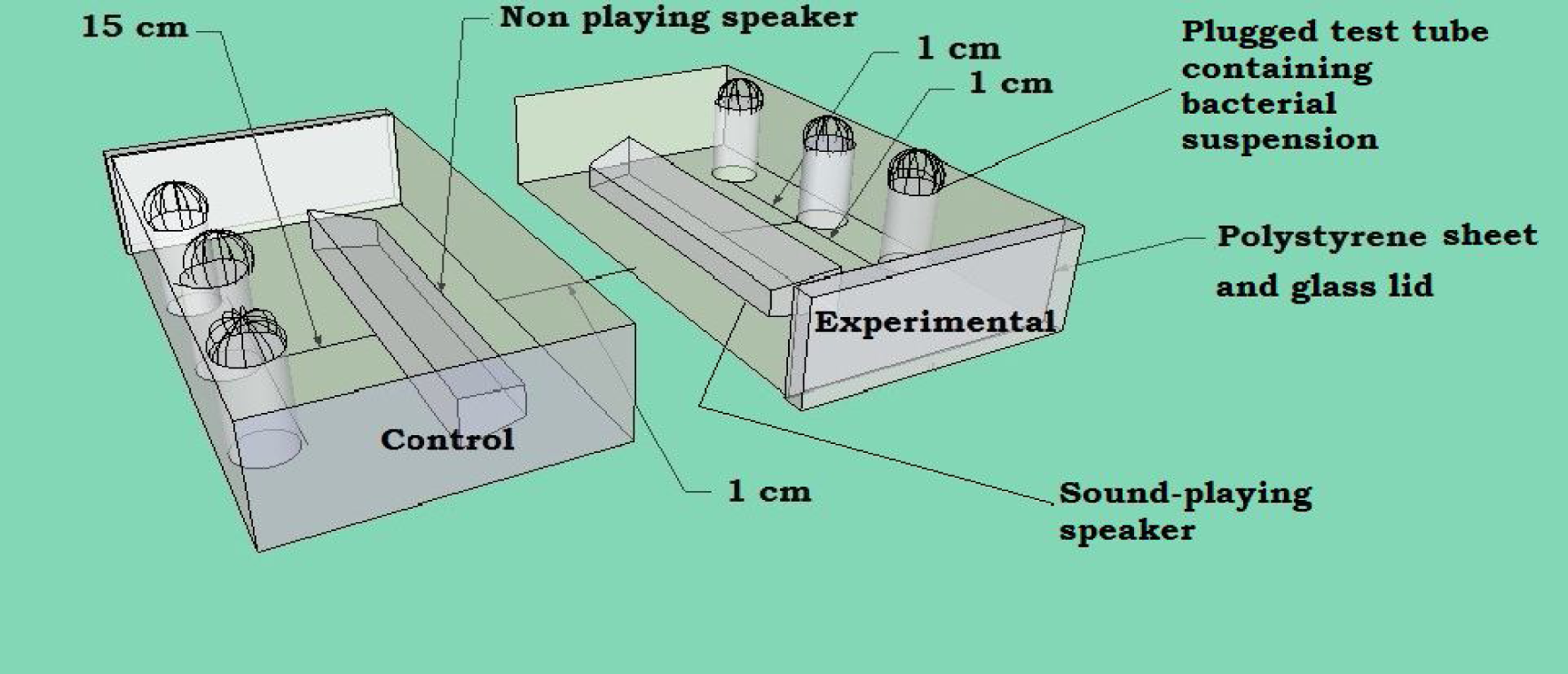
Schematic of the experimental set-up.

**Table 1:** Level of sound intensity for different frequencies Sr. No Frequency of the test sound (Hz)

| Sr. No | Frequency of the test sound (Hz) | <sup>#</sup>Intensity as measured by the sound level meter (dB) |
| --- | --- | --- |
| 1 | 100 | 80 |
| 2 | 200 | 85 |
| 3 | 300 | 90 |
| 4 | 400 | 86 |
| 5 | 500 | 84 |
| 6 | 600 | 75 |
| 7 | 700 | 82 |
| 8 | 1000 | 86 |
| 9 | 2000 | 99 |

<sup>#</sup>Volume regulating knob of the speaker was set at a constant level of 9 for all frequencies

### Growth and pigment estimation

At the end of incubation, after quantifying the cell density at 764 nm (Joshi *et al*., 2016), the culture tubes were subjected to extraction of the respective pigment, as described below.

### Prodigiosin extraction (Pradeep *et al*., 2013)

One mL of the culture broth was centrifuged at 10,000 rpm (10,600 g) for 10 min. Centrifugation was carried out at 4°C, as prodigiosin is a temperature-sensitive compound. The resulting supernatant was discarded. Remaining cell pellet was resuspended in 1 mL of acidified methanol (4 mL of HCl into 96 mL of methanol; Merck), followed by incubation in dark at room temperature for 30 min. This was followed by centrifugation at 10,000 rpm for 10 min at 4°C. Prodigiosin was obtained in the resulting supernatant; OD was taken at 535 nm. Prodigiosin unit was calculated as OD_535_/OD_764_. This parameter was calculated to nullify the effect of change in cell density on pigment production.

### Pyoverdine extraction (Unni *et al*., 2014)

Two mL of the culture broth was mixed with chloroform (Merck, Mumbai) in 2:1 proportion followed by centrifugation at 12,000 rpm (15,300 g) for 10 min. This resulted in formation of two immiscible layers. OD of the upper water-soluble phase containing yellow-green fluorescent pigment pyoverdine was measured at 405 nm. Pyoverdine unit was calculated as OD_405_/OD_764_. This parameter was calculated to nullify the effect of change in cell density on pigment production

## Results and Discussion

### Pseudomonas aeruginosa

Effect of nine different sonic frequencies was investigated on growth and pigment (pyoverdine) production of *P. aeruginosa.* All the nine sonic treatments were able to alter the growth of *P. aeruginosa* significantly, whereas pigment production was altered significantly at six different sonic treatments (Figure 2). Sound corresponding to 300 Hz, 400 Hz, and 2000 Hz was able to alter final cell density only, but not the pigment production. As production of the pigment pyoverdine in *P. aeruginosa* is regulated by quorum sensing (QS) (Stintzi *et al.*, 1998), all the frequencies capable of altering pyoverdine production may be said to have QS-modulatory effect. The maximum (52.13%) effect on QS-regulated pigment production was observed at 600 Hz.

**Figure 2.**
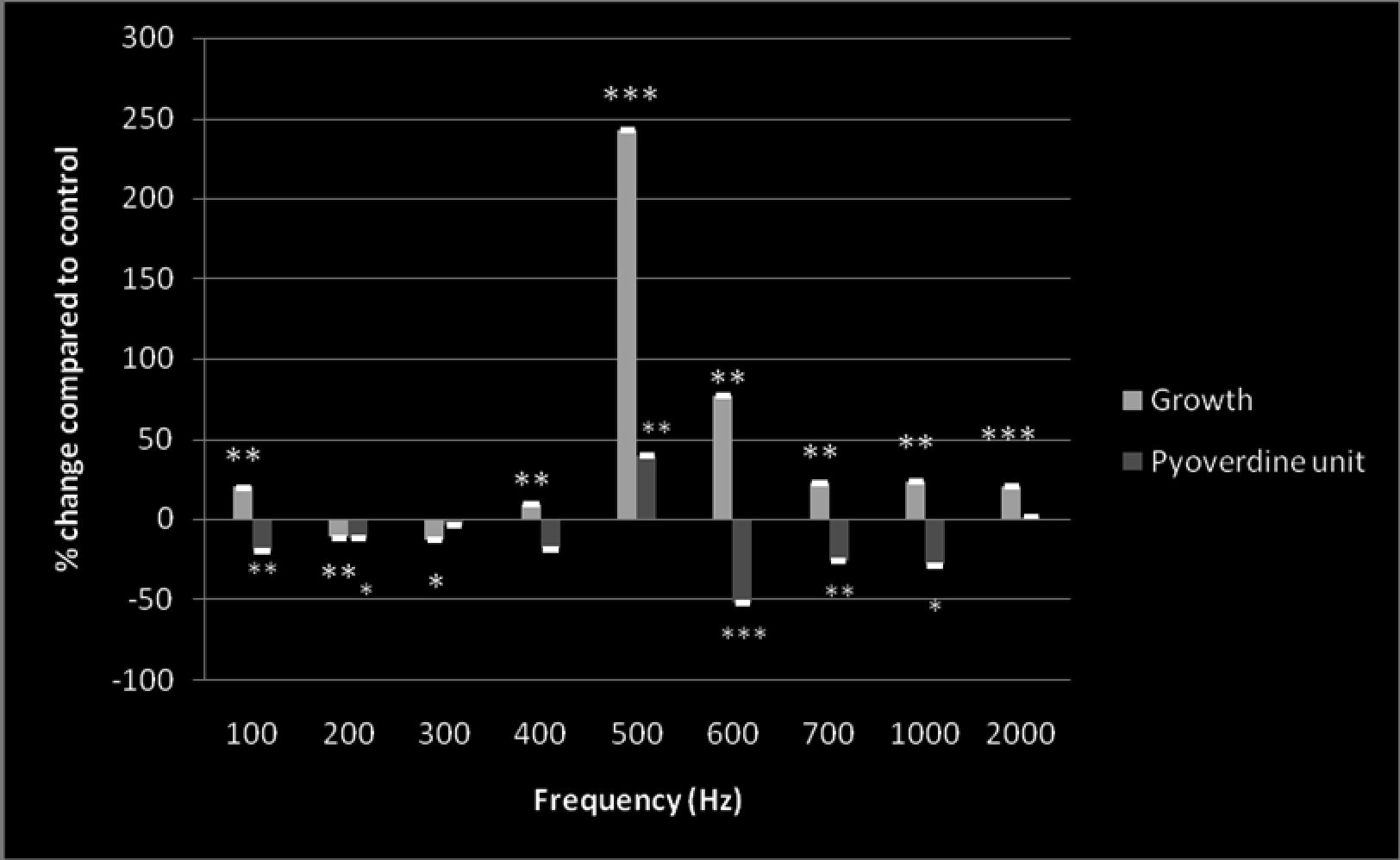
Effect of different sonic frequencies on growth and QS-regulated pyoverdine production in *P. aeruginosa*.

### Serratia marcescens

Out of the nine sonic treatments, seven (except 500 Hz and 600 Hz) were able to alter the growth of *S. marcescens* significantly (Figure 3). Production of the pigment (prodigiosin) was affected by eight different sonic frequencies; only 1000 Hz sound did not exert effect on this parameter. Thus, all the test frequencies were able to alter either one or both the test parameters (e.g. growth and pigment production).

**Figure 3.**
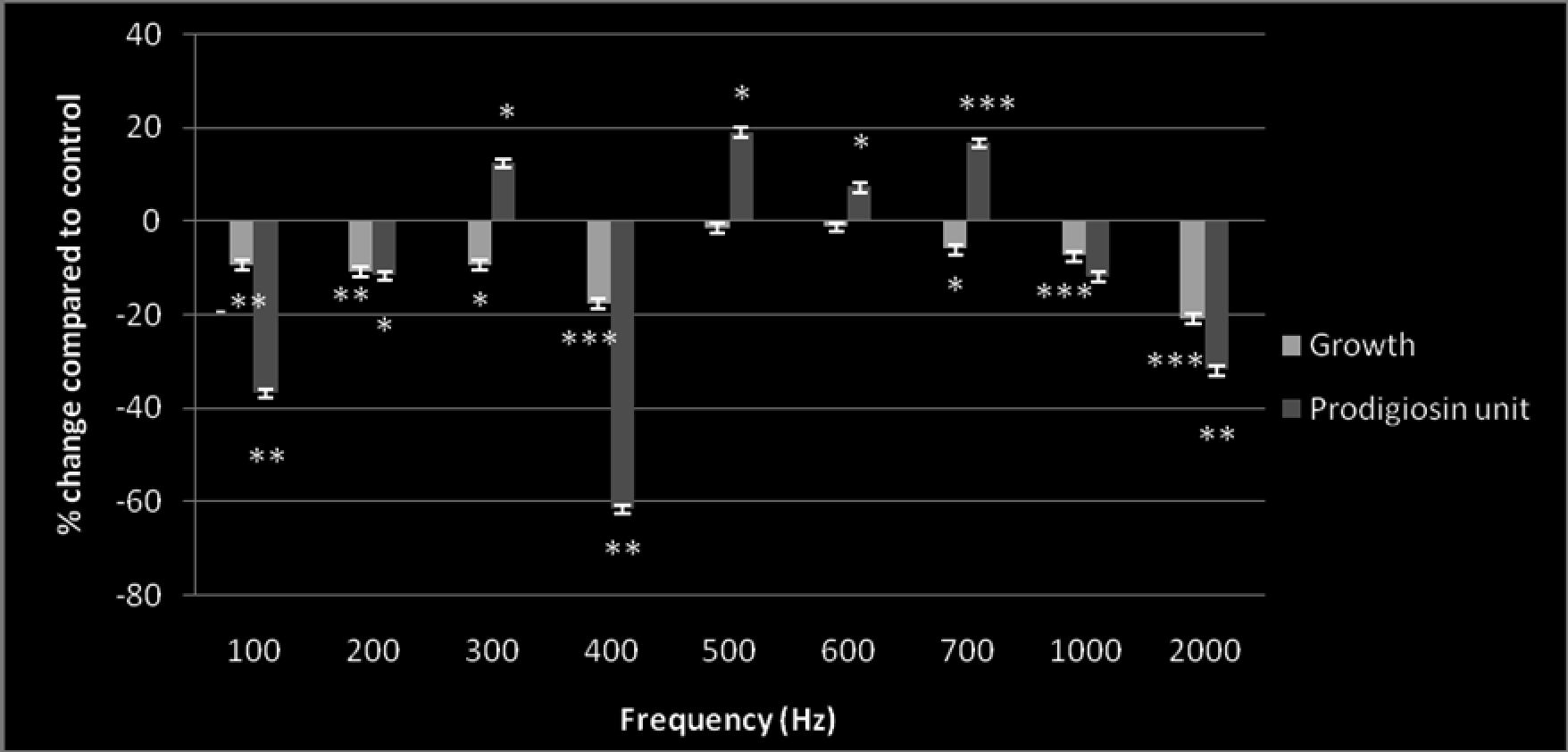
Effect of different sonic frequencies on growth and QS-regulated prodigiosin production in *S. marcescens*.

All the growth affecting sonic treatments caused *S. marcescens* to achieve a lower final cell density than control. In our previous studies (Sarvaiya and Kothari 2015a; Sarvaiya and Kothari 2015b; Shah *et al*., 2016) too, sonic stimulation was found to have an inhibitory effect on *S. marcescens* growth. 400 Hz sound affected prodigiosin production by *S. marcescens* the most; here visible prodigiosin production also started earlier in the control tube than the experimental tube. Production of the pigment prodigiosin in *S. marcescens* is known to be under regulation of QS (Morohoshi *et al*., 2007). As evident from Figure 3, except 1000 Hz, all other test frequencies had significant effect on QS-regulated prodigiosin production, and thus they can be said to possess QS modulatory effect.

Sound stimuli of 500 Hz and 600 Hz could enhance prodigiosin production (by 18.93% and 7.19 %, respectively) without affecting growth, suggesting the effect purely to be QS modulatory. Identifying the specific sonic frequencies and the differential gene expression induced by them associated with enhanced prodigiosin production can be very useful, as prodigiosin and its derivatives are of potential commercial and therapeutic value. They can find application as food colourant, proapoptotic agents for cancer treatment, potential sunscreen, etc. (Darshan and Manonmani, 2015). Prodigiosin has been reported to possess a variety of biological properties such as antimicrobial, antimalarial, anticancer, immunosuppressive, and quorum sensing inhibitory (Chaudhary et al., 2014) activities. Prodigiosin or its analogues have been considered effective biological control agents against harmful algae, have been considered cell growth regulators, and can be used as a natural dye (Lee et al., 2011).

Results of this study indicate that same frequency of sound may be responded differently by different species of bacteria. Both the bacteria used in this study are gram-negative bacteria, and hence do have some similarities with respect to their cell envelope structure, and the QS circuit possessed by them. In general gram-negative bacteria have a *luxI* / *luxR* type of AHL (acyl homoserine lactone) based QS system (Kalia, 2013). Despite these similarities, there were differences in response given by these two bacteria to same type of sonic stimulation. QS-regulated pigment production was not affected in case of *P. aeruginosa* at 300 Hz, 400 Hz, and 2,000 Hz; whereas all these three sonic frequencies were able to alter QS-regulated production of the pigment prodigiosin in *S. marcescens*. Similarly the 500 Hz and 600 Hz sound, which had only QS-modulatory effect on *S. marcescens*, affected both growth as well as pigment production in *P. aeruginosa.* In part, this difference in bacterial response perhaps may be attributed to the fact that *P. aeruginosa* has multiple QS systems (Rutherford and Bassler, 2012), and an alternative pathway may become more active in case of other pathway(s) being interfered. We studied effect of sonic stimulation only on one QS-regulated trait, however a large part of bacterial genome and a multitude of their activities are believed to be regulated by QS (Grandclement *et al*., 2015), and hence many other pathways are likely to be affected simultaneously when a sonic frequency capable of modulating bacterial QS is employed.

External sound stimulation can be considered as a type of stress for the microorganisms receiving it. Our present and previous studies, and few other researchers have already indicated that microbial growth, metabolism, and their population behavior can get altered owing to sonic stimulation. Study of the sound stimulated microbial cultures with respect to their altered gene expression can be very interesting and useful with respect to bioengineering of strains to overproduce the desired metabolite(s). For example, research into how the biology of pigment production in *S. marcescens* gets affected at the molecular level (i.e. at the level of transcriptome/ proteome) by sound stimulation can pave the way for bioengineering of prodigiosin overproducing strains. Understanding how the sonic stimulation makes bacterial culture achieve a lesser or higher cell density can enrich our knowledge regarding stress response behaviour of bacterial populations. If we can reach a stage of knowing that how a particular microbial species inside their human host will respond to a given sonic stimulation, in long run sonic waves can have therapeutic implications too. May not be in near future, but modulation of the relative abundance of different microbial populations within human microbiome, perhaps can be one of the outcomes of research in the area of microbial interaction with sonic range of sound waves.

## Acknowledgement

Authors thank Nirma Education & Research Foundation (NERF), Ahmedabad for financial and infrastructural support; Microbiology Department at M.G. Science Institute for providing the *Pseudomonas aeruginosa* culture.

